# A system for multiplexed selection of aptamers with exquisite specificity without counter-selection

**DOI:** 10.1101/2021.11.01.466780

**Authors:** Alex M. Yoshikawa, Leighton Wan, Liwei Zheng, Michael Eisenstein, H. Tom Soh

## Abstract

Aptamers have proven to be valuable tools for the detection of small molecules due to their remarkable ability to specifically discriminate between structurally similar molecules. Most aptamer selection efforts have relied on counter-selection to eliminate aptamers that exhibit unwanted cross-reactivity to interferents or structurally similar relatives to the target of interest. However, because the affinity and specificity characteristics of an aptamer library are fundamentally unknowable *a priori*, it is not possible to determine the optimal counter-selection parameters. As a result, counter-selection experiments require trial-and-error approaches that are inherently inefficient and may not result in aptamers with the best combination of affinity and specificity. In this work, we describe a high-throughput screening process for generating high-specificity aptamers to multiple targets in parallel, while also eliminating the need for counter-selection. We employ a platform based on a modified benchtop sequencer to conduct a massively-parallel aptamer screening process that enables the selection of highly-specific aptamers against multiple structurally similar molecules in a single experiment, without any counter-selection. As a demonstration, we have selected aptamers with high affinity and exquisite specificity for three structurally similar kynurenine metabolites that differ by a single hydroxyl group in a single selection experiment. This process can easily be adapted to other small-molecule analytes, and should greatly accelerate the development of aptamer reagents that achieve exquisite specificity for their target analytes.

**Significance statement:** Aptamers offer the exciting potential to discriminate between structurally similar small molecules. However, generating such highly specific aptamers has been proven challenging using the conventional process of counter-selection. In this work, we describe a high-throughput screening platform that can characterize the specificity of millions of aptamers towards a group of structurally related molecules in a single experiment and generate exquisitely specific aptamers without any counter-selection. As exemplars, we generated aptamers with high affinity and specificity towards three structurally related kynurenine metabolites using our platform. Our platform can be readily adapted to other small molecule targets and should therefore accelerate the development of aptamer reagents with exquisite specificity.

## Introduction

Aptamers are a class of oligonucleotide-based affinity reagents that have proven highly effective for the detection of small molecules in applications including molecular diagnostics (1, 2), live cell imaging (3, 4), and real-time monitoring of drugs in live animals (5, 6). This is because the aptamer selection process can be tailored to achieve desired levels of specificity, and there are even examples of aptamers that can distinguish molecules that differ by only a single methyl group (7). To increase the likelihood of generating aptamers with such high specificity, researchers typically incorporate a process known as counter-selection into their standard positive selection-based SELEX workflow. Here, the aptamer pool isolated after a given round of selection for binding to the target is incubated with one or more ‘counter-targets’ that represent common interferents or other structurally similar molecules, and aptamers that bind to the counter-targets are removed from the pool.

While conceptually straightforward, the implementation of effective counter-selection is extremely challenging because most of the key variables that govern the outcome of the selection cannot be readily measured (8–10). For example, researchers must determine the number of counter-selection steps to perform, and when to implement them within the overall selection process. Furthermore, the stringency of each individual counter-selection step must be optimized in terms of incubation time and counter-target concentration. But since the initial affinity and range of specificities for an aptamer library are fundamentally unknowable *a priori*, optimal counter-selection parameters cannot be identified in advance, and can only be determined through time-consuming trial-and-error testing. The consequences of poor counter-selection conditions can include selection of low-quality aptamers, or the outright failure of the selection (8, 9). For example, excessive counter-selection may remove desirable aptamers that offer an ideal balance of affinity and specificity but are present only at low copy numbers in the aptamer pool (11). On the other hand, inadequate counter-selection may result in cross-reactive aptamers with poor specificity. Furthermore, the process of characterizing aptamers for specificity requires significant effort due to the well-known challenges of measuring binding interactions between small molecules and aptamers (12). As such, an alternative strategy to counter-selection that can facilitate the efficient and reliable generation of high-specificity aptamers is urgently needed.

In this work, we describe a high-throughput screen that can characterize the specificity of millions of aptamers towards a group of structurally related molecules in a single experiment, and generate exquisitely specific aptamers without any counter-selection process. Our approach builds upon our previously published non-natural aptamer array (N2A2) system, in which we modified an Illumina MiSeq sequencer such that vast numbers of aptamer clusters can be generated and characterized for binding on a sequencing flow-cell (13). Our new screening procedure consists of three steps: a multiplexed enrichment step, high-throughput sequencing, and a high-throughput specificity screen. First, a conventional capture-SELEX technique is used to enrich a DNA aptamer library towards a combined pool of structurally similar molecules. Next, the enriched library is sequenced on an Illumina MiSeq, generating millions of aptamer clusters on the surface of the flow-cell. Finally, we characterize the binding of the aptamer clusters to each of the target molecules to identify highly specific aptamers. To demonstrate the utility of our platform, we characterized the specificity of an enriched aptamer library towards the tryptophan metabolite kynurenine and four structurally related kynurenine metabolites. Without any counter-selection, we were able to identify high-specificity aptamers for three of the metabolites, including multiple aptamers that can differentiate molecules differing by only a single hydroxyl group. The entire selection, sequencing, and screening process is faster than a traditional selection campaign, and typically requires ~2 weeks from the beginning of the experiment to the isolation and characterization of multiple aptamers with high specificity for closely related molecules.

## Results and Discussion

### The high-throughput specificity screening platform

Our workflow consists of three stages: multitarget enrichment, MiSeq sequencing, and a high-throughput specificity screen (**Figure 1A-C**). In the first step, the DNA aptamer library (**Supplementary Table 1**) is enriched for sequences that bind a group of small-molecule targets using a previously established capture-SELEX protocol (14, 15) (**Figure 1A**; see Supplemental Information and **Supplementary Figure 1** for more details). Briefly, a DNA library with a 30-nt variable region is hybridized to complementary 15-nt biotinylated capture strands that are coupled to a column of streptavidin-functionalized agarose resin. The column is then washed with buffer to remove unbound sequences, after which the pooled target solution is added to the column. We then collect the flow-through, which contains structure-switching aptamers that are released from the complementary capture strand upon binding to a small-molecule ligand. As proof-of-principle, we chose five molecules from the kynurenine pathway (KP)—kynurenine (Kyn), kynurenic acid (KA), 3-hydroxykynurenine (3HK), xanthurenic acid (XA), and 3-hydroxyanthranallic acid (3HA) (**Figure 1D**)—with 100 μM of each metabolite in the pooled target solution. The aptamers eluted from the column are then PCR-amplified and converted to single-stranded DNA for use in a subsequent round of screening. We performed seven rounds of this enrichment process. During the PCR amplification step, we utilized a fluorescein-labeled forward-primer, which enabled us to monitor the relative amount of the aptamer pool that survived each round of capture-SELEX (16).

**Figure 1:**
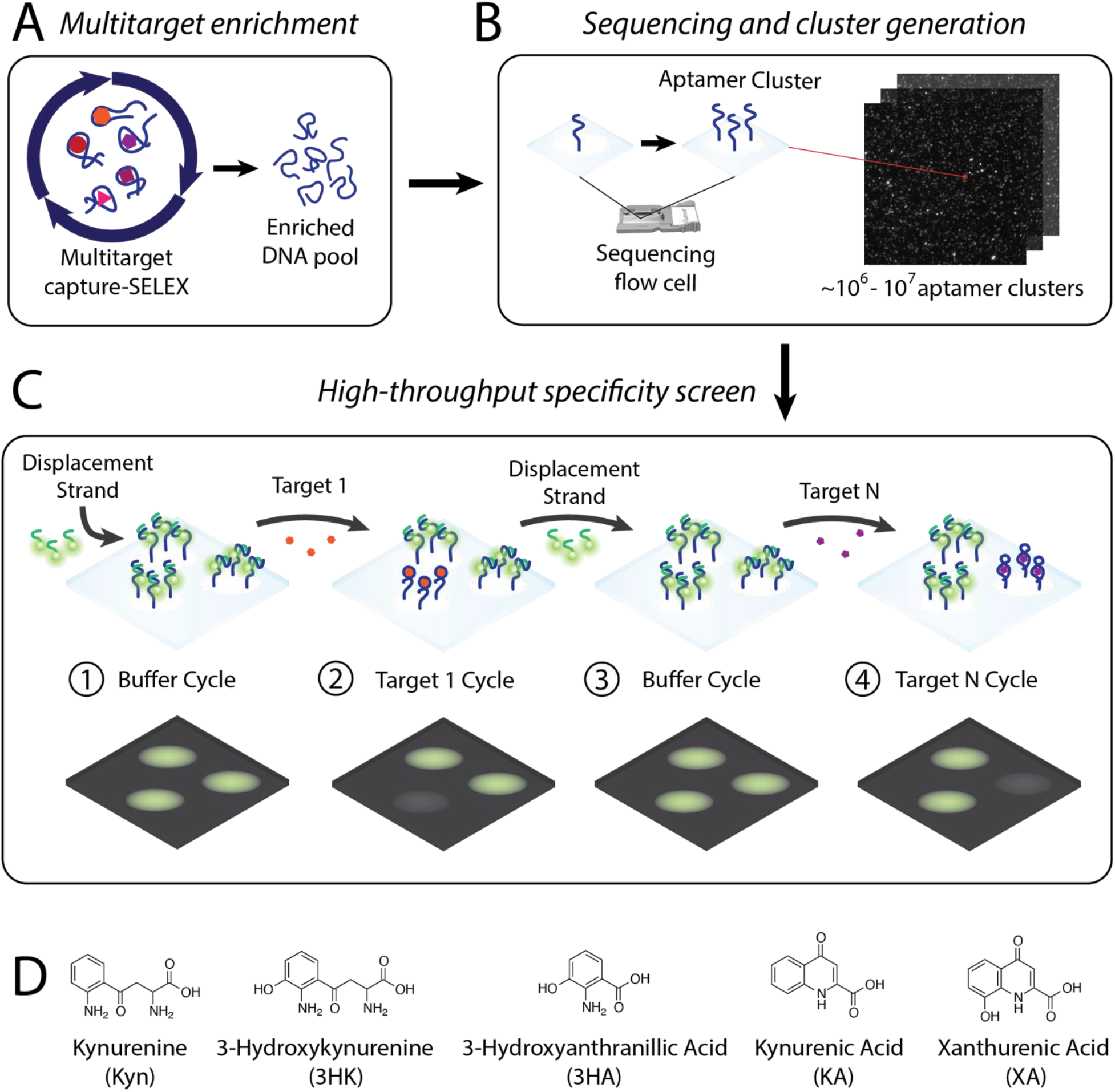
Overview of the high-throughput specificity screening platform. **A)** Multitarget aptamer enrichment. In the first step of the aptamer selection pipeline, Capture-SELEX is performed against five pooled kynurenine metabolites (Kyn, 3HK, 3HA, KA, and XA) to enrich the aptamer library for binders prior to high-throughput screening. **B)** Sequencing and cluster generation. The enriched aptamer pool is sequenced on a modified Illumina MiSeq sequencer, during which ~10^6^-10^7^ aptamer clusters are generated on the sequencing flow-cell. **C)** High-throughput specificity screen. After sequencing is complete, a ‘buffer cycle’ is conducted in which a Cy3-labeled displacement strand is annealed onto the aptamer clusters in buffer and imaged. Next, a ‘target cycle’ is conducted, in which the flow-cell is incubated with a single metabolite target for 15 minutes and then imaged. This process of buffer cycles and target cycles is repeated multiple times for each target. The flow-cell is imaged at each step, capturing the fluorescence of each cluster. **D)** Structures of the five KP metabolites used in this study.

Selections conducted against pools of structurally-similar targets tend to favor the selection of cross-reactive aptamers, and it can be exceedingly difficult to identify rare aptamers that are truly specific to individual targets. Our previously-described N2A2 system gives us a platform to identify these ‘needle in a haystack’ target-specific aptamers, in which we can measure the affinity and specificity of millions of distinct aptamer sequences on the flow-cell of a high-throughput sequencer in a single experiment (**Figure 1B-C**) (13). While the aptamer pool is being sequenced, this also results in aptamer clusters being generated on the MiSeq flow-cell. Following the first read of sequencing, the sequence following the reverse primer is removed by exploiting an EcoRI cut site within the reverse-primer region in order to minimize target interactions with Illumina adapter sequences.

The final, specificity screening stage of our assay employs a strand-displacement-based readout mechanism. Every aptamer cluster on the flow-cell is hybridized to a Cy3-labeled displacement strand, which uses the same complementary sequence as the capture strand from the selection step. And as in the enrichment step, aptamers that bind to a given target molecule undergo a conformational structure switch that results in the displacement of the fluorescently-labeled strand, resulting in a reduction in signal. We interrogate the specificity of each aptamer cluster by conducting a series of ‘buffer cycles’ followed by ‘target cycles’. In the buffer cycles, aptamer clusters are annealed with the labeled displacement strand, washed with buffer, and then imaged on the flow-cell. In the target cycles, aptamer clusters are incubated with one of the targets at a 100 μM concentration for 15 minutes, washed with buffer, and imaged again. These cycles were repeated for each of the five KP metabolites, with at least one additional replicate cycle for each target. By comparing the relative reductions in fluorescence between the five different KP targets, we can identify highly specific aptamers that do not show appreciable cross-reactivity towards the other metabolites.

### Generating a “map” of aptamer-metabolite specificity

We analyzed 2.8 million aptamer clusters during our high-throughput specificity screen of the five KP metabolites, 87% of which represented unique sequences. We processed the fluorescence intensity data to remove outliers, extremely low- or high-intensity clusters, and clusters that exhibited high variance between cycles. We measured the intensity data in terms of percent change from each buffer cycle to the subsequent target cycle, where a higher percent change indicates a greater drop in fluorescence intensity. To normalize for cycle-to-cycle differences, the percent changes for each cluster in response to each target were converted to Z-scores. This was done by subtracting the mean percent change for all clusters in a cycle from each individual percent change measurement; this value was then divided by the standard deviation of the percent change for all clusters in that cycle. Finally, we averaged these Z-scores across all replicates for each sequence-target combination. We set a cutoff Z-score of 2.576—corresponding to a 0.5% chance of false-positive binding—to identify target-binding sequences. These aptamers were further designated as target-specific if their ‘specificity ratio’—that is, the ratio of their target Z-score to the largest off-target Z-score—was greater than 3.

We then created a “map” to compare the overall binding of the aptamer pool to each pair of targets (**Figure 2**). We achieved the greatest success with 3HK, with the largest number of monospecific aptamers (902 candidates)—indeed, the majority of the 3HK aptamers were monospecific to 3HK (**Figure 2K-O**, gray diamonds). In contrast, we only identified six candidates that were monospecific for XA (**Figure 2U-Y**, orange squares) and one that was monospecific for KA (**Figure 2F-J**, red inverted triangle), and were unable to identify any monospecific aptamers for either HA or Kyn. In order to rule out the possibility that the large number of monospecific 3HK sequences were primarily mutants of a small number of aptamers that were present in the initial naïve library by chance, we grouped these sequences into families using a Levenshtein edit distance of ≤ 5. We identified 784 families among the 902 candidate clusters, indicating that the majority of the 3HK specific aptamers were not closely related and thus arose independently. These results clearly indicate that 3HK is more amenable to aptamer recognition than the other KP metabolites.

**Figure 2:**
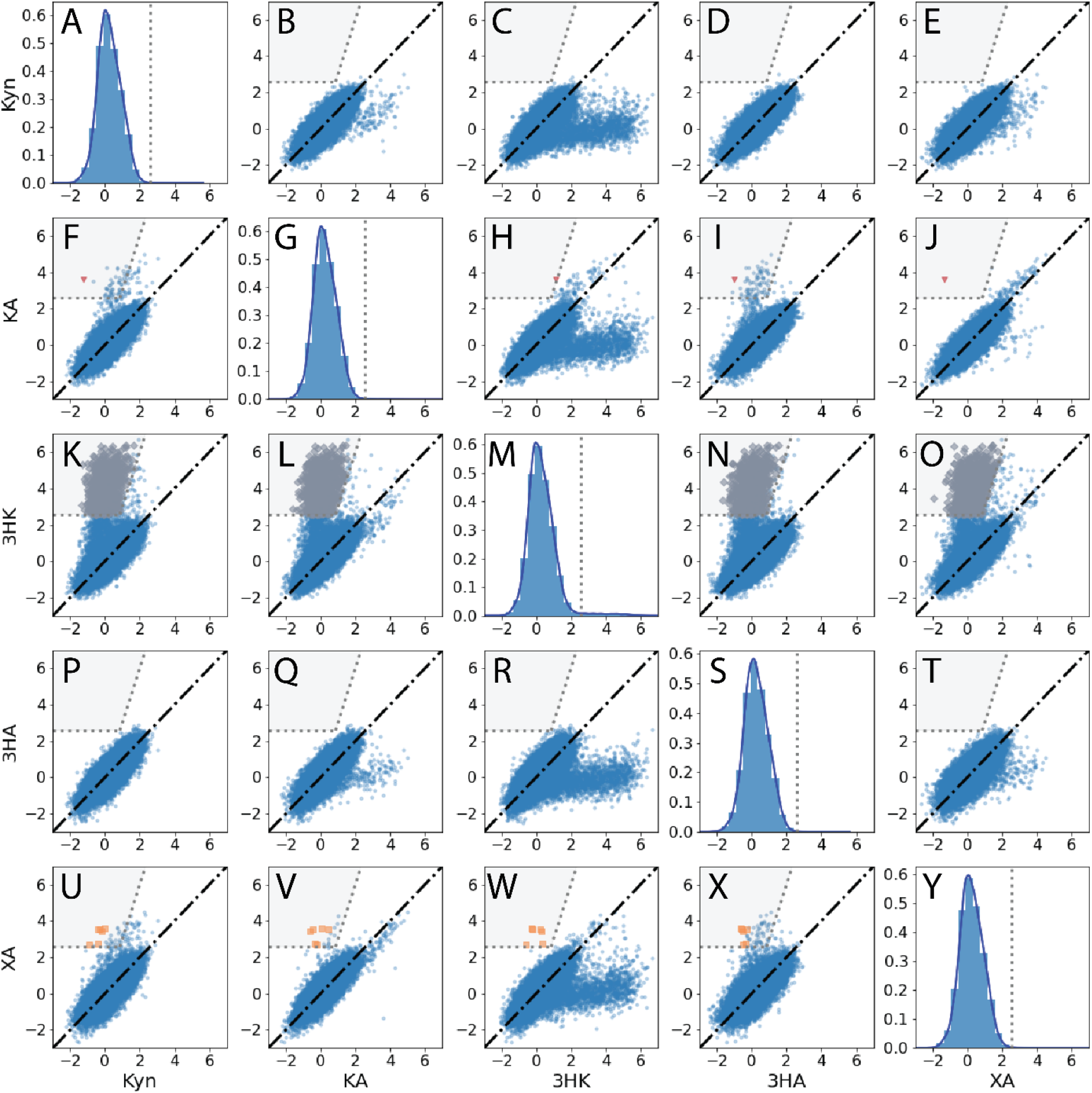
Specificity map of the aptamer pool towards the five KP metabolites. **A, G, M, S**, and **Y** show probability density functions and histograms of the Z-scores for Kyn, KA, 3HK, 3HA, and XA, respectively. The y-axis describes the probability density, indicating the likelihood that any given sequence would yield a particular Z-score. **B-E, F, H-J, K-L, N-O, P-R, T**, and **U-X** show comparisons of Z-scores between pairs of targets for all sequences with two or more replicates. Each point represents the average Z-score for a single sequence across multiple replicates. Sequences with equivalent scores for both targets fall on the black diagonal line in each plot.

In contrast, only a small proportion of the XA aptamer candidates (6/77) and KA aptamer candidates (1/90) were classified as target-specific. Most of the cross-reactive KA and XA aptamers bound both molecules with similar affinity, which was not surprising since the two metabolites only differ by a single hydroxyl group. We did note that additional monospecific aptamers for these two targets could be identified by using a less stringent specificity ratio cutoff. For example, with a specificity ratio cutoff of 2, we identified 47 more 3HK candidates and 2 more KA candidates (**Supplementary Figure 2**).

Sequences that are likely to bind to the target shown on the y-axis (Z-score ≥ 2.576) and are specific for that same target (specificity ratio ≥ 3) fall into the gray regions on the plots. Putative monospecific sequences are highlighted as red inverted triangles for KA, gray diamonds for 3HK, and orange squares for XA. No monospecific sequences were found for Kyn or 3HA.

We were not able to identify any monospecific Kyn and 3HA aptamer candidates using our stringent screening thresholds (Kyn, **Figure 2A-E**; 3HA, **Figure 2P-T**), although there were some cross-reactive aptamer candidates that displayed at least some degree of binding to each target (6 sequences for Kyn, 13 sequences for 3HA). This result was unexpected—particularly for Kyn, due to its structural similarity to 3HK, for which selection was highly successful. Although we expected many aptamers to be cross-reactive to both targets, as with XA and KA, the majority of 3HK binders did not show any cross-reactivity with Kyn (**Figure 2K**). We hypothesized that there might be extremely rare Kyn- or 3HA-binding aptamers present in the multitarget enrichment pool that were not expressed as clusters on the flow-cell during sequencing. To address this possibility, we conducted two additional rounds of selection with that pool using only 3HA and Kyn. When we repeated the specificity screen on this pool, we noted that 3HK-, XA-, and KA-specific aptamers were far less abundant, as expected. But even though some highly cross-reactive 3HA and Kyn aptamers were enriched this time around, we were still unable to identify any monospecific aptamers for these targets (**Supplementary Figure 3**). Subtle chemical features may account for the variable success rates of these screens. For example, although 3HA and 3HK share a similar aromatic moiety in their neutral forms, 3HA almost solely exists in its carboxylate form at pH 7.5 (17).

### Identification and characterization of monospecific KP metabolite aptamers

We next identified the most promising monospecific aptamer candidates for 3HK, XA, and KA from the specificity screen. We selected sequences 3HK-1 (Z-score = 4.90 for 3HK versus a maximum of 0.29 for other targets; **Figure 3A**), KA-1 (Z-score = 3.60 for KA versus a maximum of 1.11 for other targets; **Figure 3B**), and XA-1 (Z-score = 2.72 for XA versus a maximum of 0.77 for other targets; **Figure 3C**). Inspection of the flow-cell images for these aptamer clusters during the buffer and target cycles provided further evidence of their specificity and agreed with the extracted intensity values (**Supplementary Figure 4**). Interestingly, we generally observed that the highest copy-number sequence from the enrichment stage did not exhibit significant binding to any of the five targets (maximum average Z-score = −0.51) (**Supplementary Figure 5A**). This demonstrates that our screen could overcome the effects of PCR biases, contamination, or other experimental artifacts that might lead to the unwanted enrichment of low-quality sequences in other SELEX-based selection strategies.

**Figure 3:**
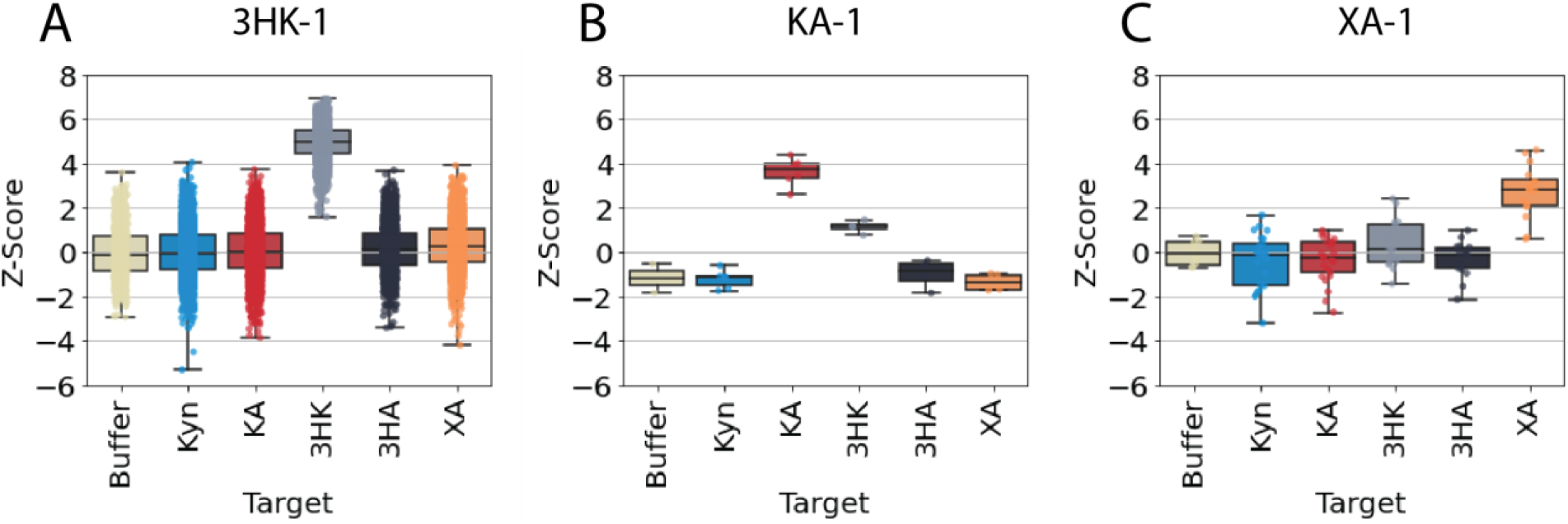
Results of the high-throughput specificity screen. Z-scores for cluster intensities for buffer and the five KP metabolites for aptamers **A)** 3HK-1, **B)** KA-1, and **C)** 3XA-1. The discrete Z-score values are overlaid for all measurements on a box plot. The middle of the box plot represents the median value and the top and bottom of the box are the upper and lower quartiles. The whiskers are the minimum and maximum values. 3HK-1, KA-1, and 3XA-1 were respectively represented by 847, 2, and 8 aptamer clusters.

While our N2A2 assay can assess each aptamer’s relative affinity for each target during the specificity screen, it cannot directly measure aptamer K_D_. We therefore utilized a previously established plate-reader assay (15), in which 3HK-1, KA-1, and XA-1 were chemically synthesized and labeled at the 5’ end with Cy3, and then combined with displacements strands tagged with a DABCYL quencher group at the 3’ end. These strands were of various lengths for each aptamer (12–14 nucleotides), where the length was selected to ensure that its K_D_ for the aptamer was ~100–500 nM, as previously recommended (see Methods for details) (15). After hybridizing the two strands, each aptamer-displacement strand complex was titrated with varying concentrations of the five different KP metabolites. Target binding results in ejection of the displacement strand, and the fluorescence intensity increases in the absence of the quencher group. The K_D_ can subsequently be derived from a quantitative equation developed for this competition assay (18). First, we characterized the binding interaction between the Cy3-labeled aptamer and DABCYL-labeled displacement strand (**Supplementary Figure 6**). Next, we measured the interaction between the aptamer-displacement strand complex and the target of interest. Finally, these measurements were used to compute the K_D_ of the aptamer for each target.

The plate-reader assay confirmed that the selected aptamers exhibited strong affinity for 3HK (3HK-1 K_D_ = 388.4 nM), KA (KA-1 K_D_ = 3.7 μM), and XA (K_D_ = 56.5 μM) (**Figure 4**). As expected, these aptamers were also remarkably specific, displaying essentially no cross-reactivity to any of the other KP metabolites at the concentrations assayed. Non-specific quenching of Cy3 was observed at high concentrations of 3HK, leading to RFU signals below baseline (**Supplementary Figure 7**). There are no previously published aptamers towards these molecules, but by way of comparison, the affinity of 3HK-1 for its target is nearly an order of magnitude higher than that of a previously published tryptophan aptamer (K_D_ ~ 2 μM (19)), which is relevant given that Kyn is a metabolite of tryptophan. Furthermore, 3HK-1’s target affinity is more than two orders of magnitude greater than the majority of previously reported DNA aptamers for amino acid targets, which typically exhibit K_D_s in the mid-micromolar to low millimolar range (20–22).

**Figure: 4:**
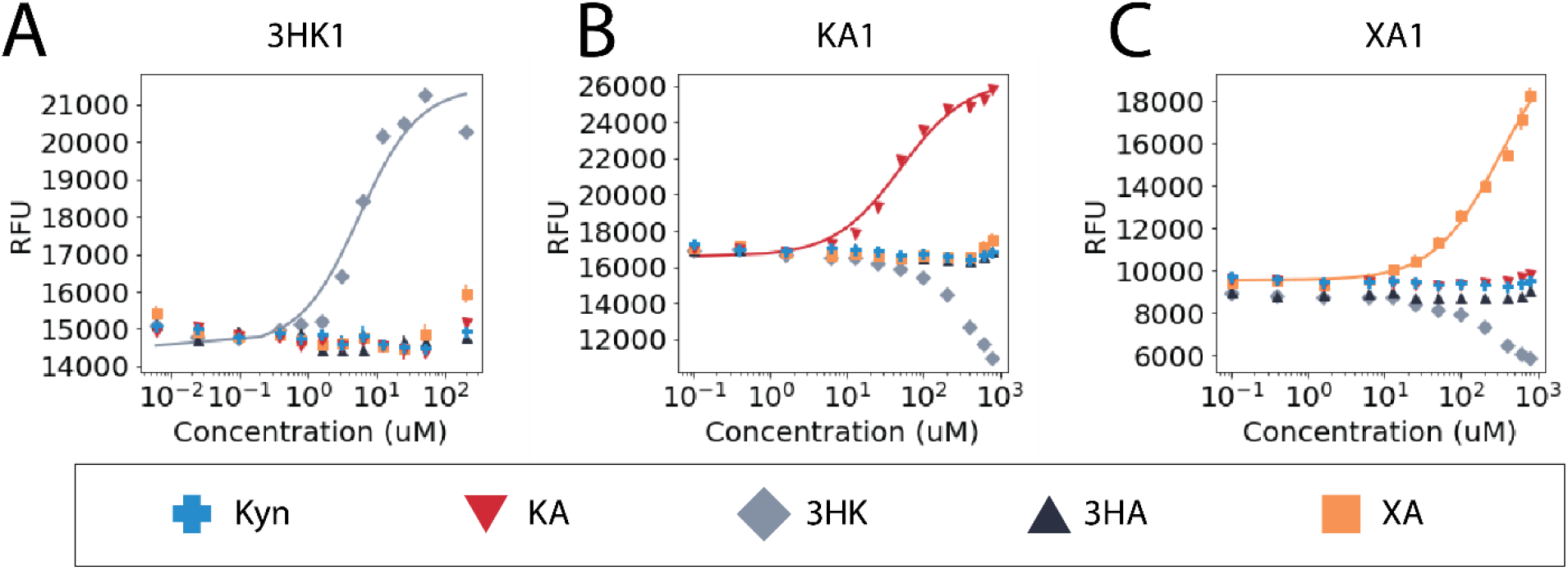
Measuring the target affinity and specificity of the 3HK-1, KA-1, and XA-1 aptamers via plate-reader assay. Binding assays for aptamers **A**) 3HK-1, **B**) KA-1, and **C**) XA-1 for the five KP metabolites. The points represent the mean of three independent experiments, and the error bars represent the standard deviation.

## Conclusion

In this work, we describe an aptamer generation pipeline that enables the efficient discovery of highly-specific aptamers for multiple structurally similar molecules in a single experiment without counter-selection. We were able to obtain specific aptamers for three of the five kynurenine metabolites included in the selection, where the selected aptamers exhibited essentially no off-target binding and can differentiate between molecules that differ by only a single hydroxyl group. The elimination of a counter-selection procedure greatly simplifies the selection workflow, bypassing the time-consuming trial-and-error process of optimization that is typically required. And as demonstrated by the poor target-binding properties we observed for the most abundant sequence from our enrichment-stage pool, the use of the N2A2 platform for screening can overcome inherent biases associated with multi-round SELEX in order to identify the top-performing aptamers in a vast sequence pool (**Supplementary Figure 5)**. Finally, our platform greatly increases the throughput of selection, such that we can identify high-specificity candidate aptamers for multiple targets in a single experiment, with the entire process of selection and characterization of individual aptamers requiring just two weeks from start to finish.

Although this study focused on monospecific aptamers, which are the most desirable for many molecular detection applications, we would like to note that this same approach can also be used to identify cross-reactive aptamers with defined specificities (**Supplementary Figure 8**). Technologies such as ‘Toggle-SELEX’ have been previously utilized to purposely select such cross-reactive aptamers that can bind proteins expressed by different two different species of animal (23). Such species cross-reactivity enables the preclinical evaluation of potentially therapeutic aptamers in animal models. Furthermore, cross-reactive aptamers that can bind promiscuously to entire families of small molecules with a shared chemical scaffold could be useful in many contexts, such as detecting illicit drugs and their metabolites, where detecting a family of small molecules is more efficient than the use of multiple highly specific assays (24). In our prior work, we have highlighted the use of N2A2 to map sequence determinants that influence target affinity (13), and we envision that data derived from this screening process could likewise uncover sequence and structural elements that inform target specificity. Given the central importance of target specificity in determining the practical utility of an affinity reagent for clinical or research applications, we believe our platform will deliver immediate value as a means for generating superior aptamers in a far more efficient manner than was possible before.

## Acknowledgements

This work was supported by the Chan-Zuckerberg Biohub, W. L Gore & Associates, the Helmsley Trust, the Biomedical Advanced Research and Development Agency (BARDA, 75A50119C00051) and the National Institutes of Health (NIH, OT2OD025342, R01GM129314-01). A.M.Y. was supported by the Stanford Bio-X Graduate Fellowship.

