## Supplemental information for "A system for multiplexed selection of aptamers with exquisite specificity without counter-selection"

**SI Materials:**

The DNA library, primers, capture strand, displacement strands, and adaptor sequencing strands were all chemically synthesized by Integrated DNA Technologies (IDT) (**Supplementary Table 1**). Fluorophore- or biotin-tagged strands were HPLC-purified, and all other sequences besides the library were purified via polyacrylamide gel electrophoresis (PAGE). GoTaq DNA polymerase was purchased from Promega (#M3005). The five kynurenine metabolites (3-hydroxyanthranilic acid, L-kynurenine, xanthurenic acid, kynurenic acid, and 3-hydroxyl-DL-kynurenine) were all ordered from Sigma-Aldrich (#148776, #K8625, #D120804, #67667, and #H1771, respectively). Pierce streptavidin agarose was ordered from Thermo Fisher Scientific (#20349) and the Micro Bio-Spin chromatography columns used during the selection were ordered from Bio-Rad (#7327204). Dynabeads MyOne Streptavidin C1 magnetic beads used for the single-strand generation of the amplified aptamer pool were purchased from Thermo Fisher Scientific (#65002). Cy3-labeled aptamer candidates were synthesized by the Stanford Protein and Nucleic Acid (PAN) Facility and cartridge-purified (**Supplementary Table 1**). The DABCYL quencher-tagged displacement strands were ordered from IDT HPLC-purified (**Supplementary Table 1**). The plate-reader assays were measured in Corning 96-well half-area black flat-bottom polystyrene microplates (Thermo Fisher Scientific, #07-201-205).

**SI Methods:**

Multi-target selection:

The multi-target selection protocol was adapted from previously published protocols14. In each round, we included five times more biotinylated capture strand than aptamer DNA in a final volume of 250 µL selection buffer (20 mM Tris-HCl, 120 mM NaCl, 5 mM KCl, 1 mM $MgCl_2$, 1 mM $CaCl_2$, and 0.01% Tween-20 in nuclease-free water). The first round used 1 nmol of DNA library, and subsequent rounds used 100-500 picomoles of single-stranded DNA. At the start of each round, the aptamer pool was annealed to the biotin-capture strand and folded by heating to 95˚C for 5 minutes and then incubating at room temperature for at least 30 minutes. Two separate first round batches were performed using 1 nmol of DNA library each and combined as an input for the second round.

250 µL of streptavidin resin was added to a column and washed five times with selection buffer, discarding the flow-throughs. This and all other washes were performed with one column equivalent (250 µL) of solution unless otherwise stated. The annealed library was then added to the column, and the library flow-through was collected and added back to the column three additional times to maximize library capture. The column with captured library was then washed with selection buffer to remove non-captured strands. Over the 7 rounds of selection, the number of washes was increased for stringency from 10 washes in rounds 1–4 to 12 washes in rounds 5 and 6 and finally 15 washes in round 7. Fifteen washes were used for rounds 8 and 9 with 3HA and Kyn. The five KP metabolites were pooled together at individual concentrations of 100 µM in

selection buffer (final volume 750 µL), and then added to the column in three 250 µL aliquots. Each column flow-through was collected separately and measured for fluorescence on the Qubit Fluorometer (Themo Fisher Scientific) from round 2 onward. After measurement, the flow-throughs from each round were pooled and concentrated in 3 kDa columns (spin at 14k RCF for 15 minutes) to a final volume ~75 µL.

The pool was then amplified for subsequent rounds. PCR reagents were added to the concentrated flow-through (20 µL of 100 µM FITC FP, 20 µL of 100 µM Biotin RP, 1 mL GoTaq Master Mix, water to a final volume of 2 mL). This mix was divided over 20 tubes and subjected to 95˚C for 2 minutes followed by cycles of 95˚C for 15 seconds, 54˚C for 15 seconds, and 72˚C for 30 seconds, followed by 72˚C for 1 min and then holding at 4˚C. We performed 10 cycles for rounds 1–3, 12 for round 4, 10 for round 5, 8 for round 6, 7 for round 7, and 8 for rounds 8–9. The amplified material was then collected and cleaned using a MinElute PCR Purification Kit (Qiagen) and quantified on a NanoDrop 2000 (Thermo Fisher Scientific) spectrophotometer and Qubit Fluorometer.

The amplicons were then converted to single-stranded DNA on beads. 300 µL of streptavidin beads were washed three times with 400 µL of 20 mM NaOH (with 15-minute incubations for each wash), and then three times with 800 µL selection buffer. The double-stranded DNA library was then added, with additional selection buffer to a total volume of 500 µL. The DNA and beads were incubated for 1 hour at room temperature and then washed twice with 800 µL selection buffer and once with 800 µL water. The supernatant was removed, and then 200 µL of 20 mM NaOH was added and incubated for 8 minutes at room temperature. 35 µL of 1M Tris-HCL and 500 µL selection buffer were then added to the supernatant, after which the solution was concentrated using 10 kDa size-exclusion columns from Amicon (#UFC501096), with two buffer exchanges in water (400 µL). Single-stranded DNA was measured using the NanoDrop and Qubit Fluorometer. This process was repeated for all rounds. The use of a fluorescein-labeled forward-primer for PCR enabled us to monitor the convergence of the aptamer pool.

<u>Preparation for high-throughput sequencing and screening:</u>

Approximately 125 ng of single-stranded DNA was added to a 300 µL PCR reaction volume with overhang adaptor RP and FP and amplified using the same protocol above for the round DNA. The adaptor product was cleaned using an Axygen AxyPrep Mag PCR Clean-up Kit (Thermo Fisher Scientific). The pool then was indexed using the Nextera XT DNA Library Preparation Kit, cleaned with the AxyPrep kit, after which approximately 135 ng of DNA was run on a 10% TBE gel. The desired DNA bands were cut out and sheared, and then mixed with 400 µL of TE and incubated overnight at room temperature. The supernatant was then separated from the gel using 0.2 micron VWR filters (spun at 14k RCF for 3 minutes). The product was then concentrated using 10 kDa size-exclusion columns, with buffer exchange in TE.

<u>High-throughput sequencing and specificity screen:</u>

The modified Illumina MiSeq sequencer previously developed by our lab for the N2A2 process was used to screen ~$10^7$ aptamer clusters10. The MiSeq first uses the aptamer pool and bridge amplification steps to generate monoclonal DNA clusters containing ~1,000 strands per cluster. In the first read, the MiSeq determines the sequence and location of every cluster on the flow-cell. Instead of performing the second read, however, these cycles are used to introduce custom reagents including the complementary strands and fluorescently-labeled target molecules. Before measuring cluster response to targets, the DNA strand beyond the reverse primer sequence is removed using a built-in EcoRI cut-site.

The high-throughput screening process measures cluster binding to each individual target over multiple cycles. Alternating buffer and target cycles are used to monitor displacement of the labeled strand upon aptamer-target binding. In buffer cycles, residual bound strands and targets are removed with 750 µL of 0.05 M NaOH plus 0.25% SDS. Next, the flow-cell is washed with 500 µL selection buffer before adding 0.2 µM displacement strand in 515 µL selection buffer. The flow-cell is then heated to 80˚C and slowly ramped down to 22 ˚C over ~30 minutes. Finally, the flow-cell is washed with 6 mL selection buffer and then imaged. The target cycles begin with a 1.25 mL buffer wash before adding the target of interest. Over a period of 42 minutes, 605 µL of 100 µM target in selection buffer is added, with 500 µL initially followed by seven periodic additions of 15 µL, with 5-minute pauses in between. The flow-cell is then washed with 6 mL selection buffer and imaged. We performed triplicate buffer/target cycle measurements for Kyn, KA, and 3HK, and duplicate buffer/target cycle measurements for 3HA and XA. Fluorescence measurements from each cycle were automatically quantified using the MiSeq software and the sequence-intensity data was linked using custom Python code (see GitHub repository for code).

<u>Identification of aptamer candidates:</u>

Screening data were first cleaned and filtered to find sequences with consistent binding behavior. Clusters with < 80 RFU were removed to account for background signal, while clusters with > 3,000 RFU were removed to account for quantification errors or imaging artifacts. We used a constant cutoff based on the coefficient of variance ($c_v = \sigma/\mu$) to remove clusters that did not have consistent RFU across cycles with the same target. The filtered data were then converted to percent changes and Z-scores to improve consistency across cycles. The percent change from buffer cycles ($\%_{change} = (I_{buffer} - I_{target})/I_{buffer}$) was first calculated, and these were shown to be normally distributed (data not shown). These percent changes were normalized to Z-scores ($Z = (\%_{change,cluster} - \mu_{cycle})/\sigma_{cycle}$) where $\mu_{cycle}$ is the mean percent change for the cycle and $\sigma_{cycle}$ is the standard deviation of the percent change for the cycle. Normalization using Z-scores was used to compensate for the gradual decrease in percent change with each subsequent cycle due to cluster degradation and other factors. Finally, the Z-scores for each target were used to calculate the average Z-score for each sequence. These were then averaged to derive a Z-score that reflects every replicate for a given sequence-target combination. This average Z-score served as a proxy for aptamer binding to the target, with higher Z-score indicating a higher likelihood of binding the target. To ensure a greater number of measurements for each sequence, only sequences present at two or more clusters were considered.

To identify sequences with relatively high binding for a target, we chose a critical value of Z = 2.576. For specificity, we assessed relative binding in terms of the ratio between the Z-score for a given target relative to the maximum Z-score for any of the off-target KP molecules:

$Ratio_{z_{aptamer}} = \frac{z_{target}}{z_{off-target}}$. A Z-score ratio cutoff of 3 was chosen to designate target-specific aptamers. Thus, for a target $t$, an aptamer was considered monospecific if both $z_t > 2.576$ and $\frac{z_t}{z_x} > 3$, where $z_x$ reflects the maximum Z-score for all other targets and buffer. Monospecific aptamer candidates with the highest copy number and Z-score were ordered from the Stanford PAN facility, so that their $K_D$ could be measured.

Characterization of aptamers via plate-reader:

We used a previously established assay for measuring the affinity of small-molecule-binding structure-switching aptamers to characterize each aptamer's $K_D$ for each target14. Overall aptamer $K_D$ is determined as the ratio of two $K_D$ values ($K_{D,1}/K_{D,2}$) for the displacement strand ($K_{D,1}$) and for binding of target to the annealed aptamer and displacement strand ($K_{D,2}$). We first tested DABCYL-tagged displacement strands of various lengths to find a length that achieves ~90% quenching of the Cy3-labeled aptamer (50 nM) with $K_D$ in the range ~100–500 nM. 50 nM aptamer was annealed with the chosen displacement strand, and then incubated with each target over a range of concentrations at a final volume of 100 µL selection buffer at room temperature. Fluorescence spectra for all samples were measured in an opaque black half-well plate at 25 °C on a Synergy H1 microplate reader (BioTeK), with filter cube (emission: 590/35, excitation: 538/63, gain: 55). Fluorescence data were fitted using the Hill equation with n = 1: $RFU = B_{max}X^n/(K_D^n + X^n) + c$. Curve-fitting was performed in Python using the 'curve_fit' function from the 'scipy' library. The final concentrations we chose based on titrations with each displacement strand (**Supplementary Figure 6**) were 800 nM 13-mer displacement strand for 3HK-1, 200 nM 14-mer displacement strand v2 for KA-1, and 400 nM 13-mer displacement strand for XA-1.

**SI Tables:**

| Name | Sequence (5' -> 3') |
|---|---|
| Library | GGCTCTCGGGACGAC-N(30)-GTCGTCCCTGAATTC |
| Biotin RP | Biotin-GAATTCAGGGACGAC |
| FITC FP | FITC-GGCTCTCGGGACGAC |
| Biotin capture strand | GTCGTCCCGAGAGCC-(18-atom hexa-ethyleneglycol linker)-biotin |
| Cy3 displacement strand | GTCGTCCCGAGAGCC-Cy3 |
| FP sequencing adaptor | TCGTCGGCAGCGTCAGATGTGTATAAGAGACAG NNNNGGCTCTCGGGACGAC |
| RP sequencing adaptor | GTCTCGTGGGCTCGGAGATGTGTATAAGAGACAG NNNNGAATTCAGGGACGAC |
| 3HK-1 aptamer | Cy3-GGCTCTCGGGACGACACGGGAAGCTTTAGGTTGAGCCATGTGCAGGTCGTCCCTG |
| XA-1 aptamer | Cy3-GGCTCTCGGGACGACCGGAGGTCTCTTTACTTTTAACCAGGTGAGGTCGTCCCTG |
| KA-1 aptamer | Cy3-GGCTCTCGGGACGACGATGGCGGTGTTTCTTTATTCGTAAATGGGGTCGTCCCTG |
| HC-1 strand | Cy3-GGCTCTCGGGACGACGATAAGTCGTTCATTCATTGTAGAGTTGTGGTCGTCCCTG |
| SK-1 aptamer | Cy3-GGCTCTCGGGACGACGGTATTGCATCTTGGAATACAGCTTTGCTAGTCGTCCCTG |
| TRI-2 aptamer | Cy3-GGCTCTCGGGACGACTTAGTGGGTCCTGATTTTTCGGGGTATCTTGGTCGTCCCTG |

| | |
|---|---|
| 13-mer displacement strand | GTCGTCCCGAGAG-DABCYL |
| 14-mer displacement strand | GTCGTCCCGAGAGC-DABCYL |
| 14-mer displacement strand v2 | TCGTCCCGAGAGCC-DABCYL |
| 14-mer displacement strand Cy3 | GTCGTCCCGAGAGC-Cy3 |

**Supplementary Table 1:** DNA sequences used during the multi-target aptamer enrichment and screening, and aptamer sequences and displacement strands used in plate-reader binding assays

**SI Figures:**

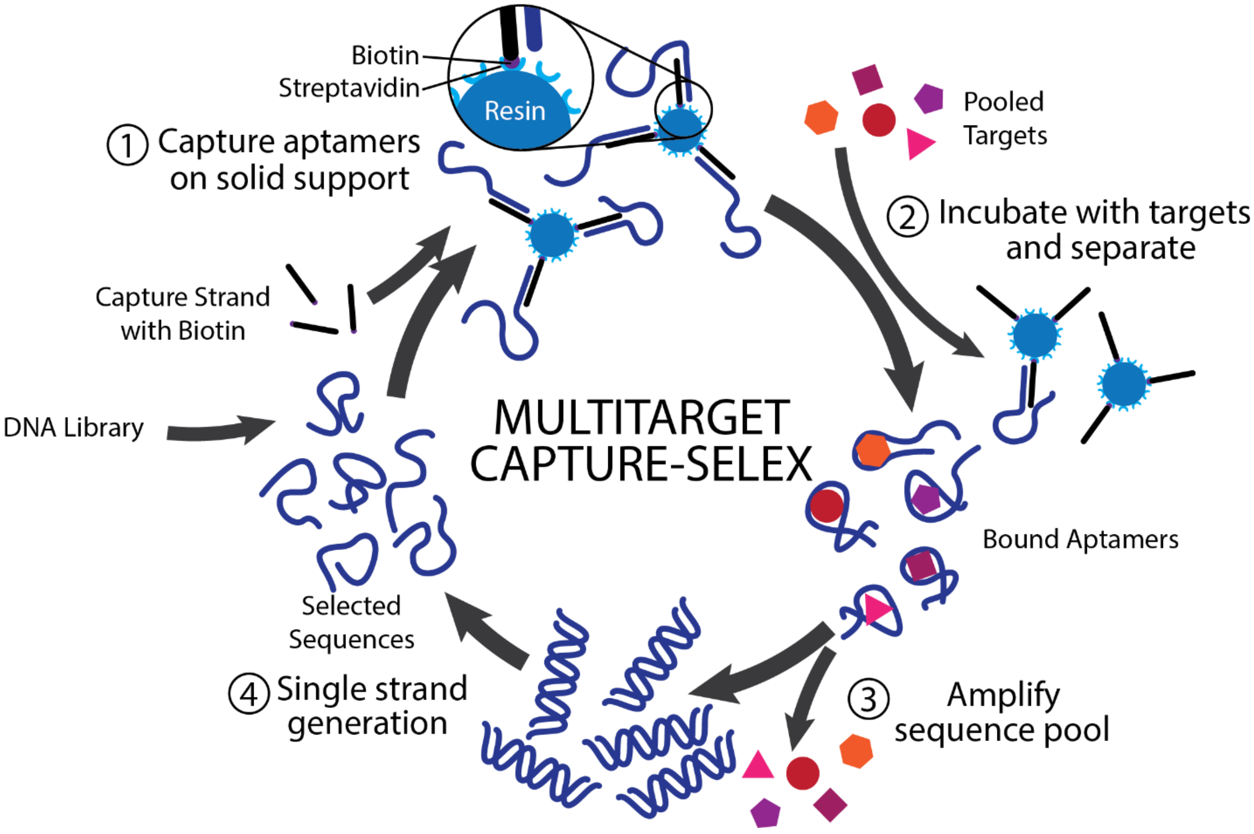


**Supplementary Figure 1:** Multitarget aptamer Capture-SELEX scheme. In the first step of the multitarget aptamer selection, the DNA library is incubated with a biotinylated capture strand and then captured onto a streptavidin functionalized solid support. In the next step, the aptamers are incubated with the pooled metabolites and the eluted structure-switching aptamers are collected. These aptamers are then PCR amplified and converted to single-stranded DNA. The selection process is then repeated several times before moving onto the high-throughput specificity screen

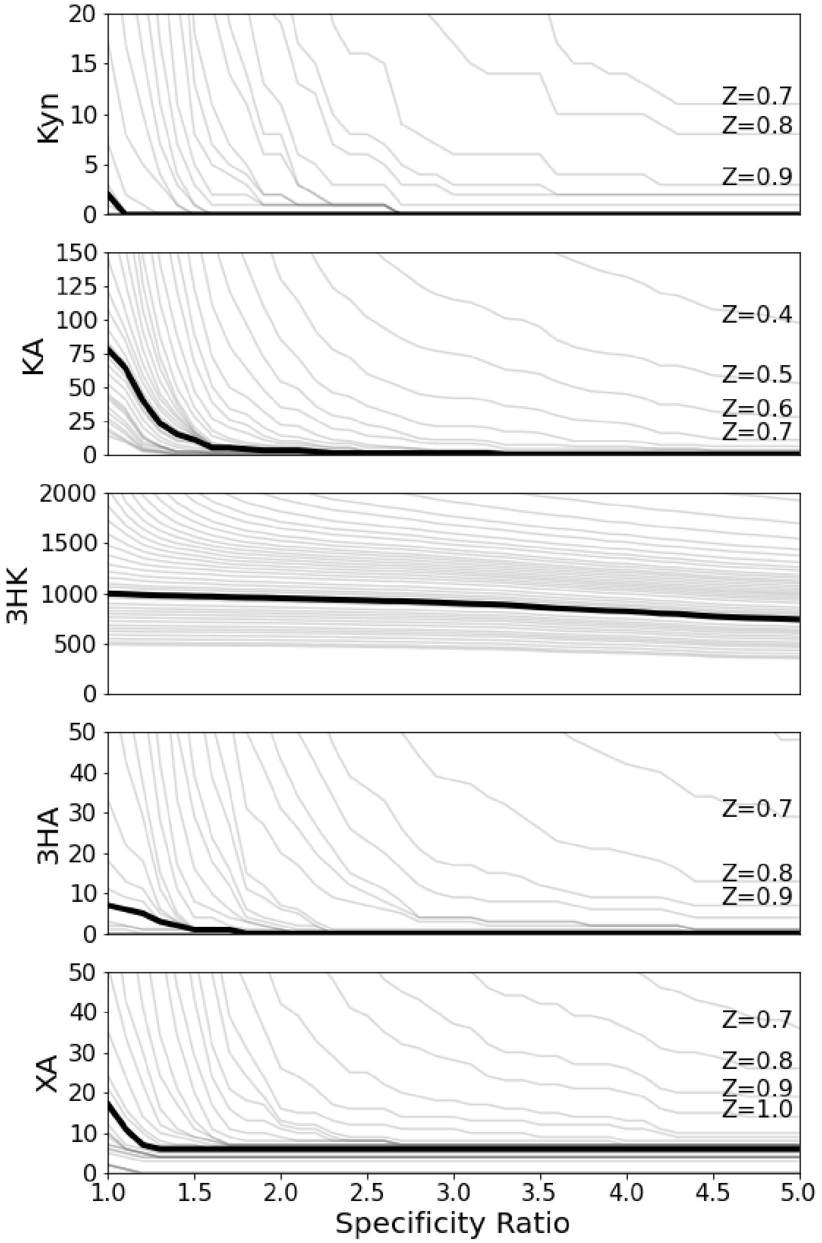


**Supplementary Figure 2:** Number of monospecific aptamer candidates for each target (y-axis) at various Z-score cutoffs (individual lines depict same Z-score cutoff), and specificity ratio cutoff values (x-axis). The bolded line represents the Z-score cutoff value of 2.576 that we ultimately used.

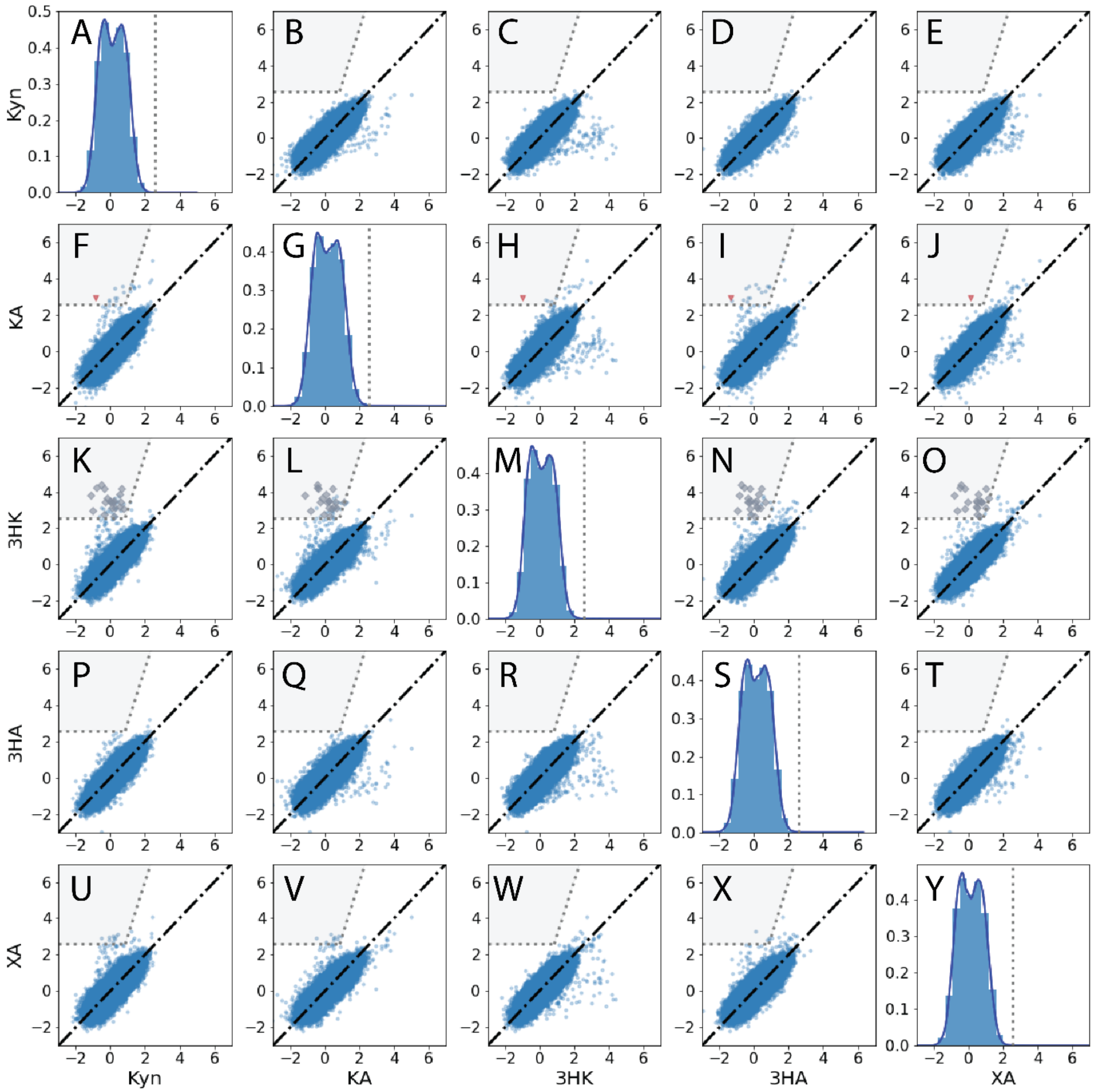


**Supplementary Figure 3:** Specificity map of the aptamer pool towards the five KP metabolites after two additional rounds of enrichment with Kyn and 3HA. **A, G, M, S,** and **Y** show probability density functions and histograms of the Z-scores for Kyn, KA, 3HK, 3HA, and XA, respectively. The y-axis describes the probability density, indicating the likelihood that any given sequence would yield a particular Z-score. **B-E, F, H-J, K-L, N-O, P-R, T,** and **U-X** show comparison of Z-scores between pairs of targets for all sequences with two or more replicates. Each point represents the average Z-score for a single sequence. Sequences with equivalent scores for both targets fall on the black diagonal line in each plot. Sequences that likely bind to a given target ($Z \geq 2.576$) and are specific (specificity ratio $\geq 3$) to the target listed on the y-axis fall into the gray region. Putative monospecific sequences are highlighted as red inverted triangles for KA and gray diamonds for 3HK. No monospecific sequences were found for Kyn, 3HA, and XA.

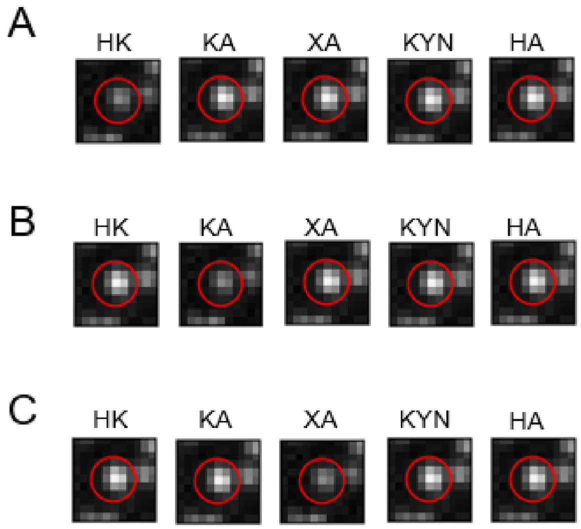


**Supplementary Figure 4:** Images of a representative cluster on the MiSeq flow-cell for each of the three monospecific aptamers. Cluster images are provided in the presence of each of the five kynurenine metabolites.

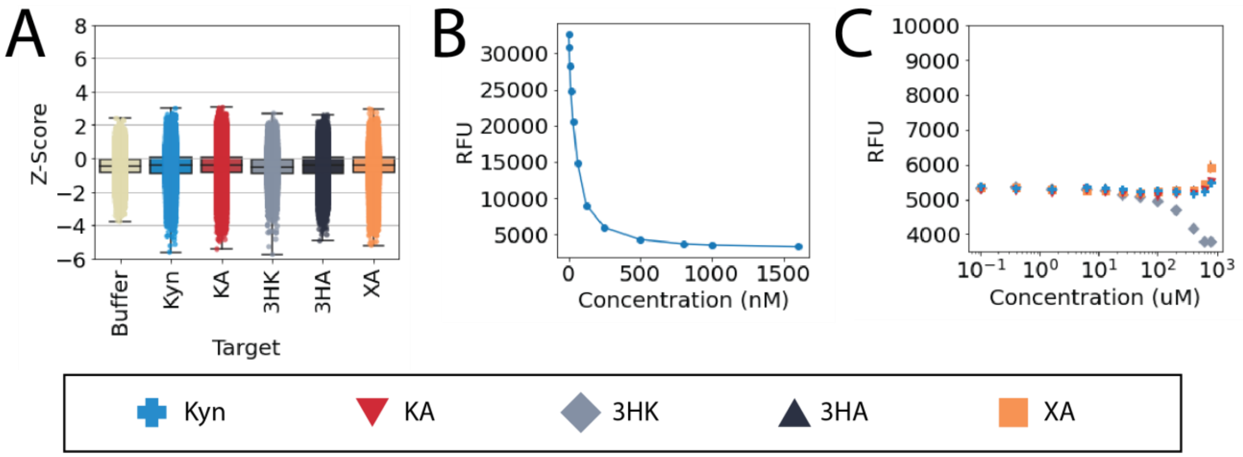


**Supplementary Figure 5:** Highest copy number sequence (HC-1) from the multi-target enrichment stage. **A)** High-throughput screening data for 108,334 clusters of HC-1 shows low Z-scores for buffer and all targets. **B)** Binding of HC-1 with 14-mer displacement strand v2 (N=1). **C)** Binding assay for HC-1 against the five KP metabolites. The points represent the mean of three independent experiments, and the error bars represent the standard deviation. Experiments were done with 400 nM 14-mer displacement strand v2 and 50 nM Cy3-labeled HC-1 aptamer.

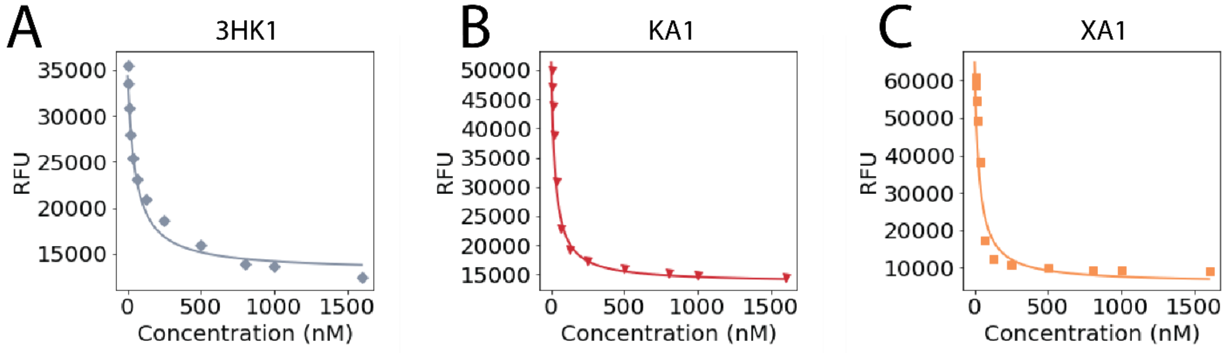


**Supplementary Figure 6:** Binding assays for aptamers **A**) 3HK-1, **B**) KA-1, and **C**) XA-1 against the displacement strands.

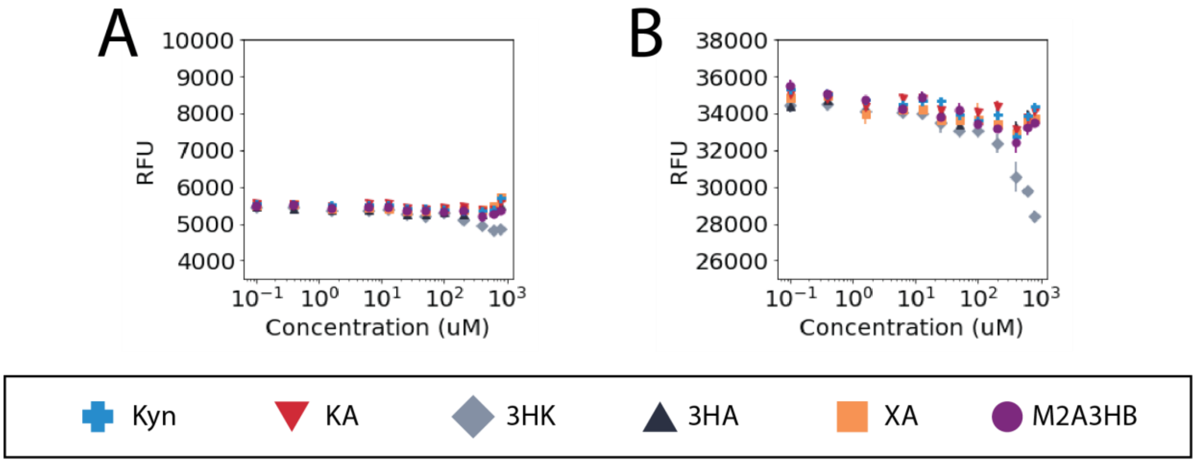


**Supplementary Figure 7:** Examination of non-specific target effects on **A)** 5 nM or **B)** 50 nM Cy3-labeled 14-mer displacement strand DNA.

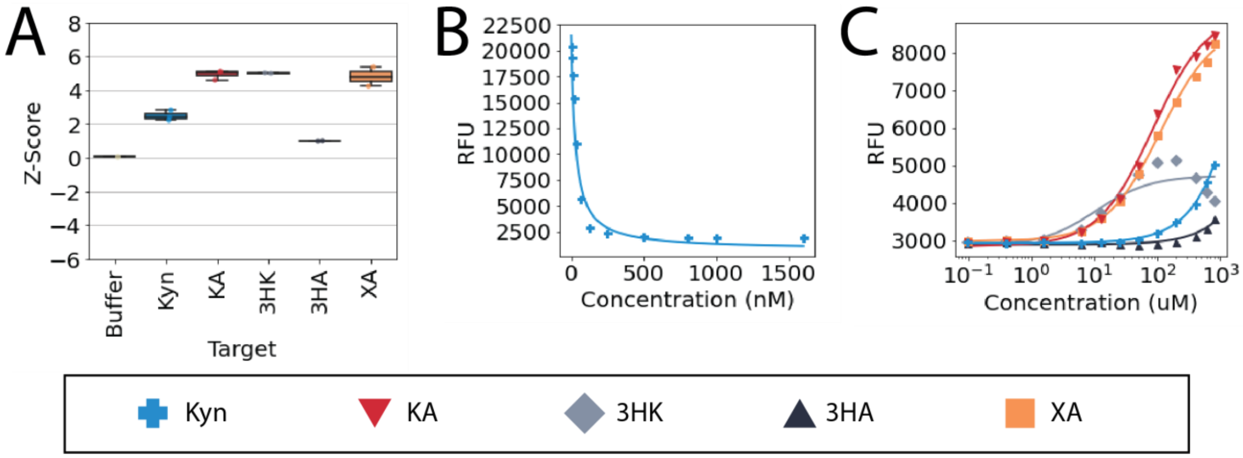


**Supplementary Figure 8:** Plate reader binding assay for cross-reactive aptamer SK-1. **A)** Flow-cell screening data for N=1 SK-1 cluster showing cross-reactive binding. **B)** Binding interaction between Cy3-labeled SK-1 aptamer and the 14-mer displacement strand v2 (N=1). **C)** Binding assay of SK-1 against the five metabolites with final displacement strand concentration of 125 nM (N = 3).
